# Introducing py_emra: the Python module for Ensemble Modeling Robustness Analysis

**DOI:** 10.1101/065177

**Authors:** Matthew K Theisen

## Abstract

The py_emra module implements the Ensemble Modeling Robustness Analysis (EMRA) algorithm in Python. Previous implementations in MATLAB have been used to gain useful insights into metabolic dynamics and metabolic engineering strategies. The py_emra package performs favorably in comparison to MATLAB. Potential extensions and improvements are discussed.

## Introduction

Kinetic simulation of metabolic systems requires realistic rate equations and parameters. Additionally, dynamic stability has been raised as an important issue in literature reports. Ensemble Modeling Robustness Analysis is a simulation framework addressing these issues.

The py_emra module is available at: https://github.com/theis188/py_emra.

## MATLAB origins

The Ensemble Modeling Robustness Analysis (EMRA) algorithm has most recently been implemented in MATLAB [1–3]. Non-standardized MATLAB implementations of the EMRA algorithm are available at https://github.com/theis188/Rivera-theisen-paper-code and https://github.com/theis188/theisen-ploscomp-bio. While MATLAB is a convenient and powerful computational environment, the open-source availability and larger user base of Python make it a favorable language for the application of the EMRA framework.

The EMRA framework uses an ensemble of kinetic models to represent a metabolic system. While a single model may be preferable, it is usually not possible to obtain exact values for enzyme kinetic parameters. Additionally, the EMRA framework uses parameter-domain integration to determine the effect of changes in enzyme amount, with potential computational savings.

## Algorithm inputs

i. **Stoichiometric matrix** The stoichiometric matrix is a matrix of connectivity for the reaction network of the system. The stoichiometric matrix should include exchange reactions (reactions to stand for input/output from the system). Each column represents a reaction and each row represents a metabolite. The matrix is sparse (most entries are 0). Negative entries (-1,-2, etc) represent substrate and positive entries represent products.
ii. **Enzyme reversibilities** Reversibilities are represented as 0 (irreversible), and 1 (reversible). The reversibility of the enzyme helps determine what rate law should be used.
iii. **Reference flux** The reference flux represents a specific steady state (dX/dt=0) of the system we’re interested in. In some cases, the reference flux is completely determined by the steady state requirement. In other cases, determination of a relevant steady state constitutes a non-trivial portion of research work.

## Algorithm Features

i. **Automated rate law definition** The algorithm uses network information (i.e. number of products, number of reactants and reversibility) to determine a suitable rate law for each reaction. There are currently 10 reaction types available which support 0,1 or 2 products for each reaction with either reversible or irreversible kinetics.
ii. **Automated parameter value generation** The parameter sets are generated by constraining the parameter sets to the reference state and dynamic stability. The normalized affinity (K_m_) and equilibrium (K_eq_) parameters are sampled from the random uniform range [1,10]. The normalized affinity parameter is equal to K_m_/X_ss_ where K _m_is the actual affinity parameter and X_ss_ is the steady state metabolite concentration. Thus, the [1,10] range indicates an unsaturated enzyme. This represents a relatively flexible system, in which rates change dynamically with metabolite concentrations and has proven computationally useful, independent of physiological accuracy. [2]
iii. **Metabolite concentration normalization** The forms of the rate laws used for the algorithm allow for normalization of the metabolite concentration. Each place where the metabolite concentration appears in the rate law, it is divided by an affinity parameter (i.e. a K_m_ value). The metabolite concentration and affinity parameter are normalized to the steady state metabolite concentration X_ss_. The fact that normalized values (affinity parameters and metabolite concentrations) ensures the normalization factor is always canceled out. The normalization simplifies the determination of metabolite concentration, since all values can be set to 1. [2]

## Algorithm performance

The algorithm uses the numpy, scipy and sympy packages extensively in its core functions. It also currently uses the xlrd package for import of the model inputs in .xls format.

The integrates in parameter-space, constrained to steady state. In MATLAB implementations, the MATLAB integrator EventFunctions are used to terminate integrations where a metabolite concentration becomes negative, or the system becomes unstable (time-domain Jacobian becomes singular).

The integrators available in the numpy package do not have EventFunction capability. The simple workaround is to perform the integration in steps. This allows the stability to be checked at each point. However, more steps slows the algorithm considerably. As the number of steps approaches 1, the algorithm improves dramatically over the MATLAB implementation. However, without a number of steps, the system will not be able to determine exactly where a system bifurcates. A custom integrator with EventFunction-like support could improve the performance of the integration and obviate the need to choose a StepNo parameter.

The system was benchmarked on a simple metabolic system to show the effect of the StepNo parameter on performance. The system was tested on a simple system with an analytical solution and the numpy integrator showed good performance, within 0.01% of the actual solution, even with StepNo=1. By default the ‘lsoda’ integrator from scipy.integrate is used.

**Table 1.** Comparison of Python and MATLAB m

| Version | Benchmark time (s) |
| --- | --- |
| MATLAB | 167.3 |
| Python (StepNo = 25) | 206.3 |
| Python (StepNo = 5) | 62.6 |
| Python (StepNo = 1) | 28.8 |

## Extensions and applications

The modular rate law of Liebermeister [4] allows for arbitrary numbers of products and reactants, and for different types of substrate level regulation in rate laws. These rate laws have been previously implemented in a modular fashion in the MATLAB versions of the EMRA framework. Addition of these to py_emra could enhance its capabilities.

Currently, import of reaction list is through a .xls file. The format is specified in a provided template. In principle, py_emra could be integrated with SBML-ready python packages like python-libsbml.

